# Protein CREATE enables closed-loop design of *de novo* synthetic protein binders

**DOI:** 10.1101/2024.12.20.629847

**Authors:** Alec Lourenço, Arjuna Subramanian, Ryan Spencer, Michael Anaya, Jiapei Miao, William Fu, Eric Chow, Matt Thomson

## Abstract

Proteins have proven to be useful agents in a variety of fields, from serving as potent therapeutics to enabling complex catalysis for chemical manufacture. However, they remain difficult to design and are instead typically selected for using extensive screens or directed evolution. Recent developments in protein large language models have enabled fast generation of diverse protein sequences in unexplored regions of protein space predicted to fold into varied structures, bind relevant targets, and catalyze novel reactions. Nevertheless, we lack methods to characterize these proteins experimentally at scale and update generative models based on those results. We describe Protein CREATE (Computational Redesign via an Experiment-Augmented Training Engine), an integrated computational and experimental pipeline that incorporates an experimental workflow leveraging next generation sequencing and phage display with single-molecule readouts to collect vast amounts of quantitative binding data for updating protein large language models. We use Protein CREATE to generate and assay thousands of designed binders to IL-7 receptor *α* and insulin receptor with parallel positive and negative selections to identify on-target binders. We discover not only individual novel binders but also features of ligand-receptor binding, including preservation of the IL7R*α* - ligand hydrophobic interface specifically and existence of multiple approaches to contact the insulin receptor. We also demonstrate the importance of structural features, such as the lack of unpaired cysteine residues, toward design fidelity and find computational pre-screening metrics, such as interchain predicted TM scoring (iPTM), while useful, are imperfect predictors as they neither guarantee experimental binding nor rule it out. We use the data collected from Protein CREATE to score designs from the initial generative models. Globally, Protein CREATE will power future closed-loop design-build-test cycles to enable fine-grained design of protein binders.

## Introduction

The ability to produce proteins with defined functional properties has enormous potential for areas including biologics, where binding tightly to a receptor target with minimal off-target effects and long half life are desired, and industrial processes, where bio-based manufacturing promises to replace syntheses that are complex, inefficient, or currently require hazardous components. The full potential of protein engineering has yet to be unleashed, however, because many applications require specific control over a protein property. For instance, taste and olfaction modulators require both high on and off rates so that the sensory receptors can quickly respond to the stimulus and reset. Other applications require optimization of multiple properties. For instance, therapeutics should have low immunogenicity and off-target binding in addition to their primary function of binding a given target. Current techniques lack the precision to generate proteins with the full set of desired properties *ab initio* and must utilize costly and time-consuming screens to optimize candidates with no guarantee of success.

Traditional protein design approaches, such as directed evolution, rely on making small mutations to a starting sequence to locally optimize protein fitness on a task. This is typically a natural protein with weak, but non-zero, activity on the property being selected for. Unfortunately, natural protein sequences are far from ideal starting points for directed evolution, as most natural proteins are marginally stable. Mutations that would be tolerated by a stable protein can destabilize a marginally stable protein to the point of non functionality[1]. This makes fitness landscapes more difficult to traverse, increasing the risk of failure. Recombination methods, such as DNA shuffling, can somewhat alleviate this problem, [2], but are still limited by the number of functional parent sequences being tested[3]. Computationally generated protein starting sequences can be less prone to be marginally stable and better able to take on multiple desired properties. For instance, Neo2/15, a computationally designed IL-2/IL-15 mimetic, not only binds to the IL-2R*βγ* dimer more than IL-2, but also has enhanced solubility when expressed in *E. coli* and thermostability when disulfide stapled[4]. Achieving this feat took multiple rounds of screening to test and optimize multiple designs due to low success rates of computational designs from physics-based tools such as Rosetta [5], which rely on heuristics to make the computational complexity of modeling protein designs tractable.

Recent “physics-free” approaches using artificial intelligence leverage natural protein sequences and structures as training data to dramatically improve the success rate for *de novo* designed proteins, sometimes with success rates upwards of 10% without the need for experimental optimization [6, 7]. The use of natural protein data is a strength and weakness of these approaches, as the algorithm has proven struggles to extrapolate its predictions to unnatural, designed sequences. While the use of metrics derived from these algorithms, such as the Alphafold interface predicted template modeling (iPTM) score, has proven to be a useful computational binary predictor of binding, it is far from sufficient to guarantee experimental binding success [8]. Leading practitioners emphasize the drive to achieve “one design, one binder”[7] via algorithmic improvements while limiting the need for experimental search. While this may lead to short-term success, history suggests that leveraging search - in this case, development of fast and cheap experimental data collection and integration - will be more a effective design approach[9]. Forward progress will require approaches that can collect large experimental data sets on designed binders and then applied to improve quality of AI models.

We present Protein CREATE (Computational redesign via an experiment-augmented training engine), a protein design framework powered by an experimental platform that collects quantitative binding data on thousands of designs against multiple targets in parallel over the course of 3 days (Figure 1a). We show Protein CREATE enables hypothesis-driven design by not only identifying individual novel protein binders but also relevant structural and biochemical features in libraries of engineered binders.

**Figure 1:**
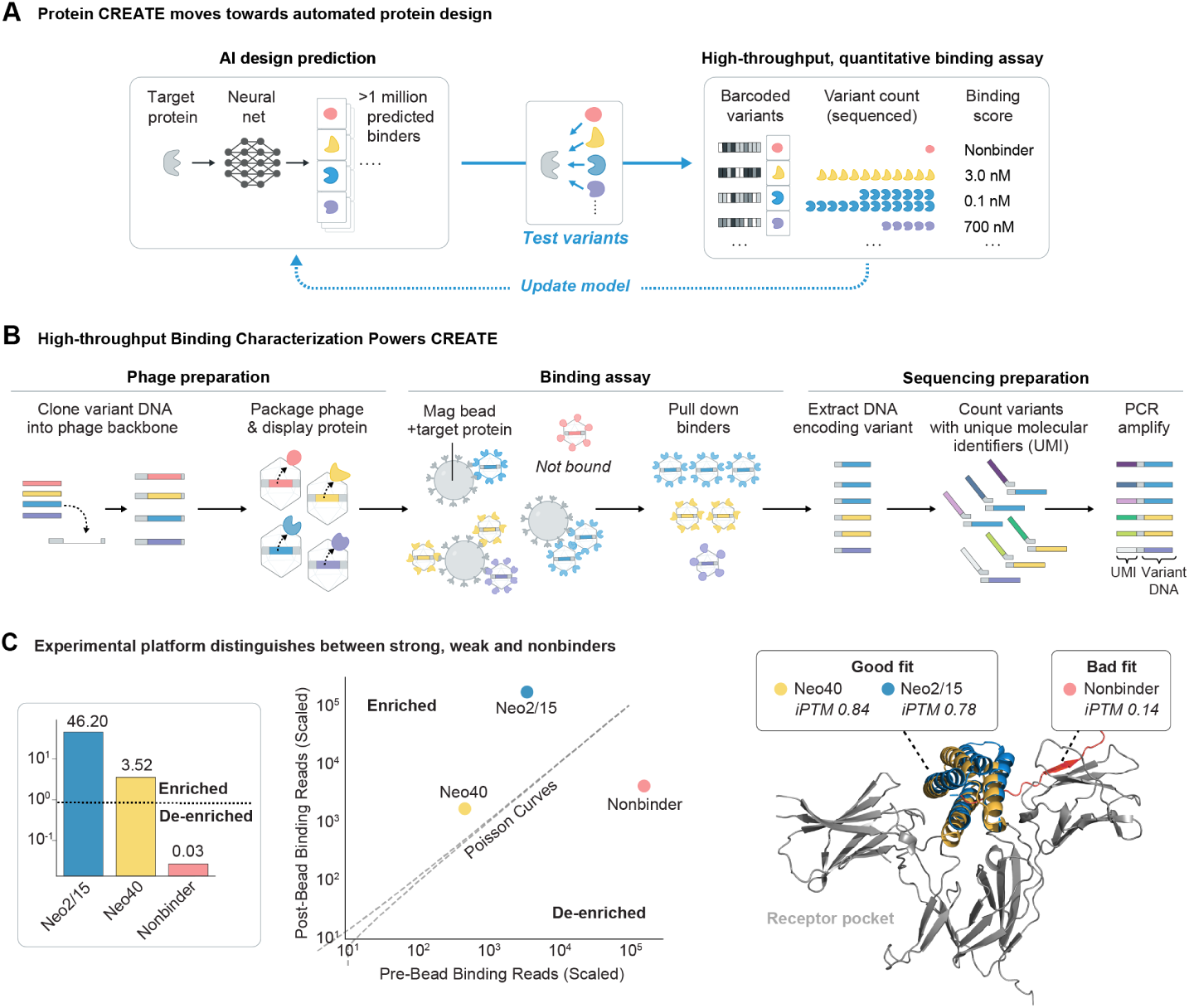
Protein CREATE is an integrated protein design platform powered by a quantitative experimental platform that ranks protein binders based on sequencing data. **a)** Protein CREATE combines the ability of AI-based design strategies to generate diverse binding candidates with an experimental assay that is high-throughput enough to rank thousands to millions of variants. **b)** Overview of the binding assay. Variants are first cloned into linear T7 bacteriophage backbones, packaged, and allowed to infect helper E. coli to display the library on their capsid surfaces. They are then purified and introduced to beads decorated with binding targets. Phages are pulled down, washed, and their DNA is extracted. To compute enrichment over the base pool, phage samples both before and after binding to target undergo sequencing preparation steps. A unique molecular identifier (UMI) is added to the DNA to identify which DNA molecules originated from a single parent phage, which reduces measurement noise and enables molecular counting, in turn making pseudo-Kd estimation possible. After an additional amplification step where Illumina adaptors are added, the amplicon library is sequenced. **c)** Previously characterized IL2 mimetics were displayed on the T7 bacteriophage capsid and were allowed to bind beads coated with IL2R*βγ*. Enrichment, as defined by the number of normalized unique reads post-binding divided by the number of normalized unique reads pre-binding, leads to clear, separable scores for previously characterized strong, weak, and off-target binders.

## Results

### Protein CREATE Accurately Ranks Protein Binding Strengths

Protein CREATE uses a phage-based “binding by sequencing” assay to quantify the binding affinity of protein libraries at scale. DNA libraries are cloned into phage backbones before phage propagation and their encoded proteins displayed on the phage capsid. The expressed library is then allowed to bind to target-immobilized beads, which is washed to remove non-binding variants. Bound phage have their genomes extracted and labeled with a unique molecular identifier (UMI), a unique barcode which facilitates molecular counting of individual variants when sequencing. Counts of bound phage are compared to their counts in the initial library to assess the binding strengths of each variant (Figure 1b).

To evaluate Protein CREATE’s ability to score protein binding strength, we tested designed IL-2R*βγ* binders whose binding strength has been previously characterized using surface plasmon resonance [4] (SPR) and a non-binding protein in our assay. As expected, phage-expressed binders were enriched while phage expressing the non-binder were de-enriched (Figure 1c). Furthermore, the degree of enrichment of the stronger binder, Neo2/15 (*K*_*D*_ = 18.8 nM), is roughly 13-fold over the weaker binder, Neo40 (*K*_*D*_ = 260 nM), in close agreement with their respective binding strengths.

### Protein CREATE Discovers Novel Binders

To demonstrate the utility of Protein CREATE, we designed and screened candidate binders against IL-7RA, a cell-surface receptor with therapeutic relevance and for which previous groups have had high success in designing binders [10, 11, 6]. As is the case for many cytokine receptors, developing both agonists and antagonists for IL7R*α* has therapeutic justification, as receptor activation can lead to T-cell proliferation [12] while inhibition can help prevent autoimmune attacks [13]. To probe the relevant interactions at the binding interface, we used a design strategy we termed “context dependent inverse folding” (Figure 2b) in which we used the structure of a known synthetic IL7R*α* antagonist in complex with IL7R*α* as a template for an inverse folding model, ESM-IF [14]. Only the binder sequence was masked, providing both structural and interface constraints, and the model is prompted to predict a novel binder sequence. We used this strategy to design 42 sequentially diverse binding candidates and screened them for binding using Protein CREATE (Figure 2c).

**Figure 2:**
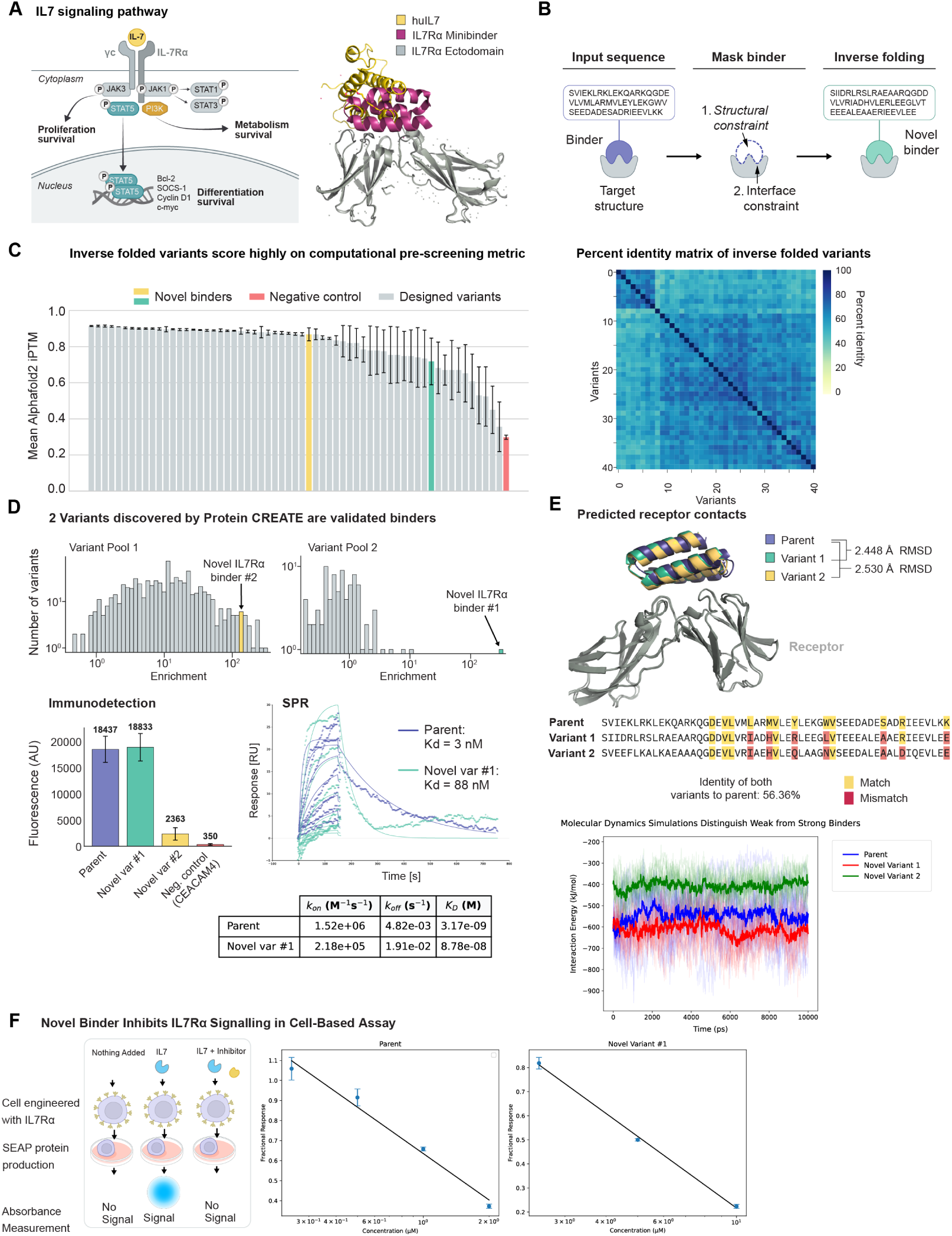
Protein CREATE discovers novel IL7R*α* binders designed using context-based inverse folding. **a)** Natural IL7 signals by binding first to IL7R*α*, then recruiting the common *γ*-chain before activating the JAK/STAT and PI3K/AKT pathways. A previously engineered mini binder binds to the IL7R*α* in a different orientation than native IL7, acting as a competitive inhibitor. **b)** Context-based inverse folding relies on masking binder sequence, but not structural, information of a known binder-target pair and unmasking binder residues using a sequence-to-sequence transformer. **c)** By using context-based inverse folding with PDB 7OPB as the binder-target pair, we designed 42 variants with high predicted binding to IL7R*α* using AlphaFold2 iPTM scores as a heuristic. **d)** We screened a subset of the variants along with a pool of off-target controls using Protein CREATE and discovered one high-affinity binder and one low-affinity binder to IL7R*α*. Follow-up experiments using a dual antibody detection strategy indicate binding of the enriched variant in an orthogonal assay when purified protein is allowed to bind to IL7R*α* immobilized on Bioplex beads. Additionally, binding for the parent binder and the stronger novel binder was characterized using surface plasmon resonance. **e)** The predicted structures for both novel variants are closely, but not perfectly, aligned to the parent structure when co-folded in AlphaFold3. The novel context-dependent inverse folded variant is predicted to fold into a closely aligned structure as the parent sequence while being only 56.4% identical. Amino acid differences are distributed across the protein sequence, including within residues predicted to contact IL7R*α* (highlighted). We performed molecular dynamics simulations on Alphafold3 predicted structures for the parent and two variants using GROMACS. Average (darker shade) and individual trajectories (lighter shades) are plotted. **f)** We screened the parent and novel variant for IL7R*α* inhibition in an engineered HEK293 cell line. Cells were incubated with 17 pM human IL7 and varying concentrations of each inhibitor. Cells express secreted embryonic alkaline phosphatase (SEAP) proportional to the degree of receptor activation, which is measured via conversion of a chromogenic substrate.

We expressed and purified two novel binders with sequence identity less than 60% to the parent sequence. Both variants were confirmed to show binding activity using a combination of SPR and an immunoassay using the Bioplex system. The strongest binding variant was shown to have a dissociation constant within two orders of magnitude of the parent and to inhibit IL7 signaling in an *in vitro* assay with engineered human cells (Figures 2d, 2f). Molecular dynamics simulations corroborate our Bioplex-based immunodetection assay, as parent and novel variant 1 have lower interaction energies compared to novel variant 2, which has a higher interaction energy and exhibits less binding experimentally (Figure 2e). We also confirmed that the predicted structures of both variants align closely with the parent when co-folded with the receptor using Alphafold 3[15]. Given the structural similarity, we inferred that the identified variants and parent should share receptor contact positions. Interestingly, about only half of the amino acid identities at these positions were preserved (Figure 2e).

### Protein CREATE Reveals Permissiveness of the IL7R*α* Binder-Receptor Interface

Given the ability of Protein CREATE to screen many candidate binders in parallel, we sought to ask what similarities enriched variants share when compared to the base pool of designs. As context-based inverse folding preserves the 3d structure of binder variants compared to the parent, including interface contacts, we analyzed the identity of the residues at positions predicted to constitute the receptor-binder interface as predicted by RING [16]. Surprisingly, a majority of predicted receptor contacts are not preserved (Figure 3a), even among enriched variants. IL7R*α* is known to have a moderately hydrophobic interface, which led us to hypothesize that more nonpolar contacts will be preserved. We compared amino acid identities at positions identified by RING in variants enriched in the assay to those in the base design pool and found that, indeed, most (3/4) are nonpolar amino acids (Figure 3b).

**Figure 3:**
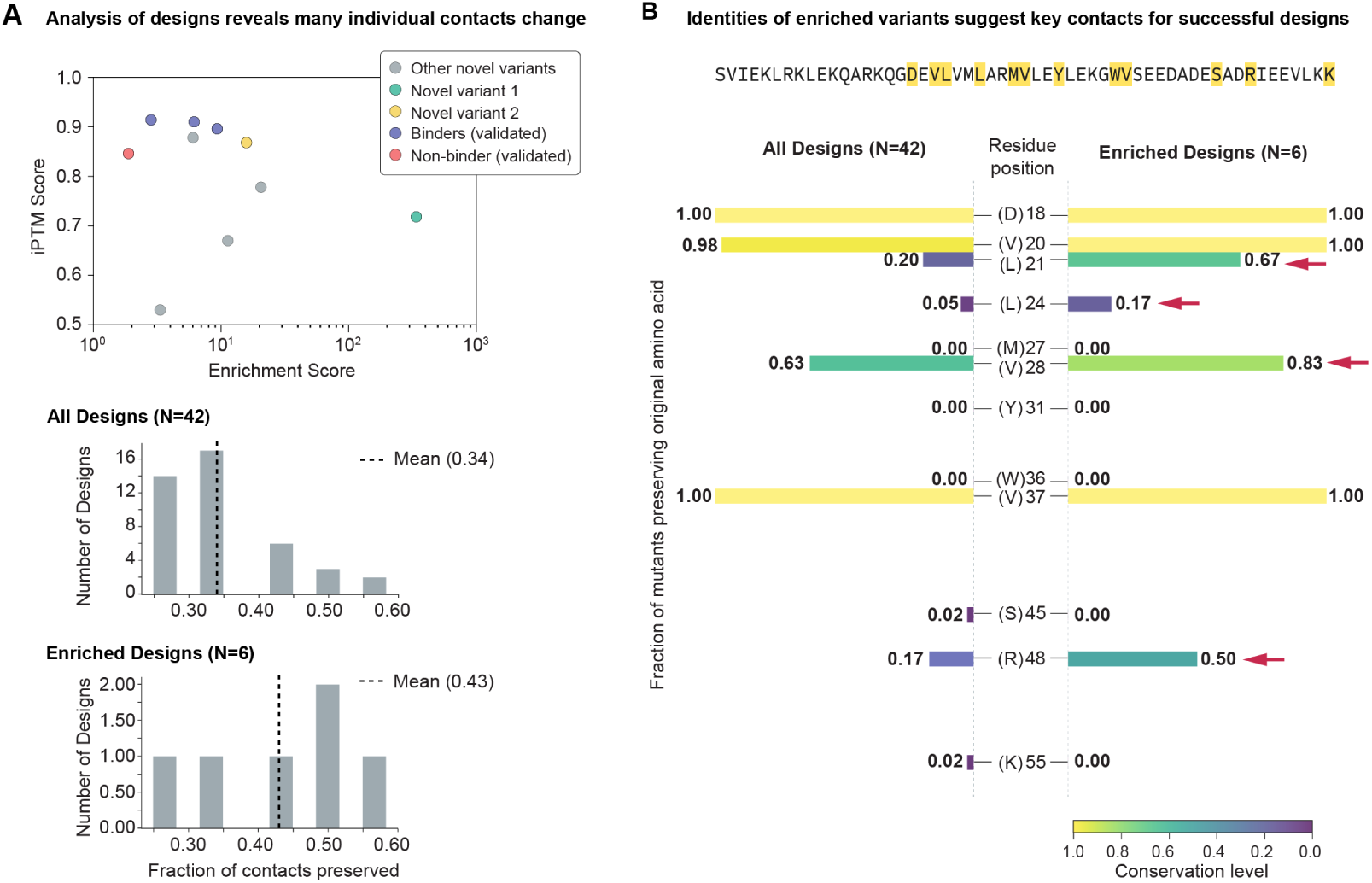
Protein CREATE discovers key receptor-binder contacts for IL7R*α*. **a)** (Top) Six variants designed via inverse folding are enriched in the Protein CREATE assay when compared with a known non-binder and several known binders previously identified [10]. Although serving as a pre-screen for designs, the interface predicted TM score (iPTM score) of the designed variants does not correlate with their relative enrichments. (Bottom) Context-dependent inverse folding does not preserve most receptor contacts, although relatively more contacts are preserved among enriched designs. **b)** Individual analysis of receptor-binder contacts indicates that context-dependent inverse folding preserves certain residues in a majority or all designs. Residue positions with more preservation among enriched designs are designated with arrows.

Additionally, CREATE allows discrimination of residues positions where the original amino acid need not be preserved (S45 and K55) from those that maintain the interface, even when the design methodology rarely preserves the amino acid identity in the base position (L21 and R48).

### Protein CREATE Interrogates the Importance of Biochemical Features in Insulin Mimetics

Insulin and insulin-like peptides are a highly conserved class of proteins within eukaryotes, with evidence for ancestral insulin-like peptides dating back at least to the advent of unicellular eukaryotes [17]. Human insulins and its homologs consist of 3 disulfide bonds - 2 inter-chain and 1 intra-chain (Figure 4a). Binding contacts between insulin and its receptor are highly conserved across vertebrate species [18], suggesting that these may be essential for binding. However, insulin-like peptides, such as *Drosophila melanogaster* insulin-like peptide 5 (DILP5), refute this assumption. While DILP5 shares structural characteristics with human insulin, such as both consisting of an A and B chain stapled together by 3 disulfide bonds, DILP5 only retains 2 out of the 8 receptor contacts yet binds to human insulin receptor with a *K*_*D*_ of 60 nM [19].

**Figure 4:**
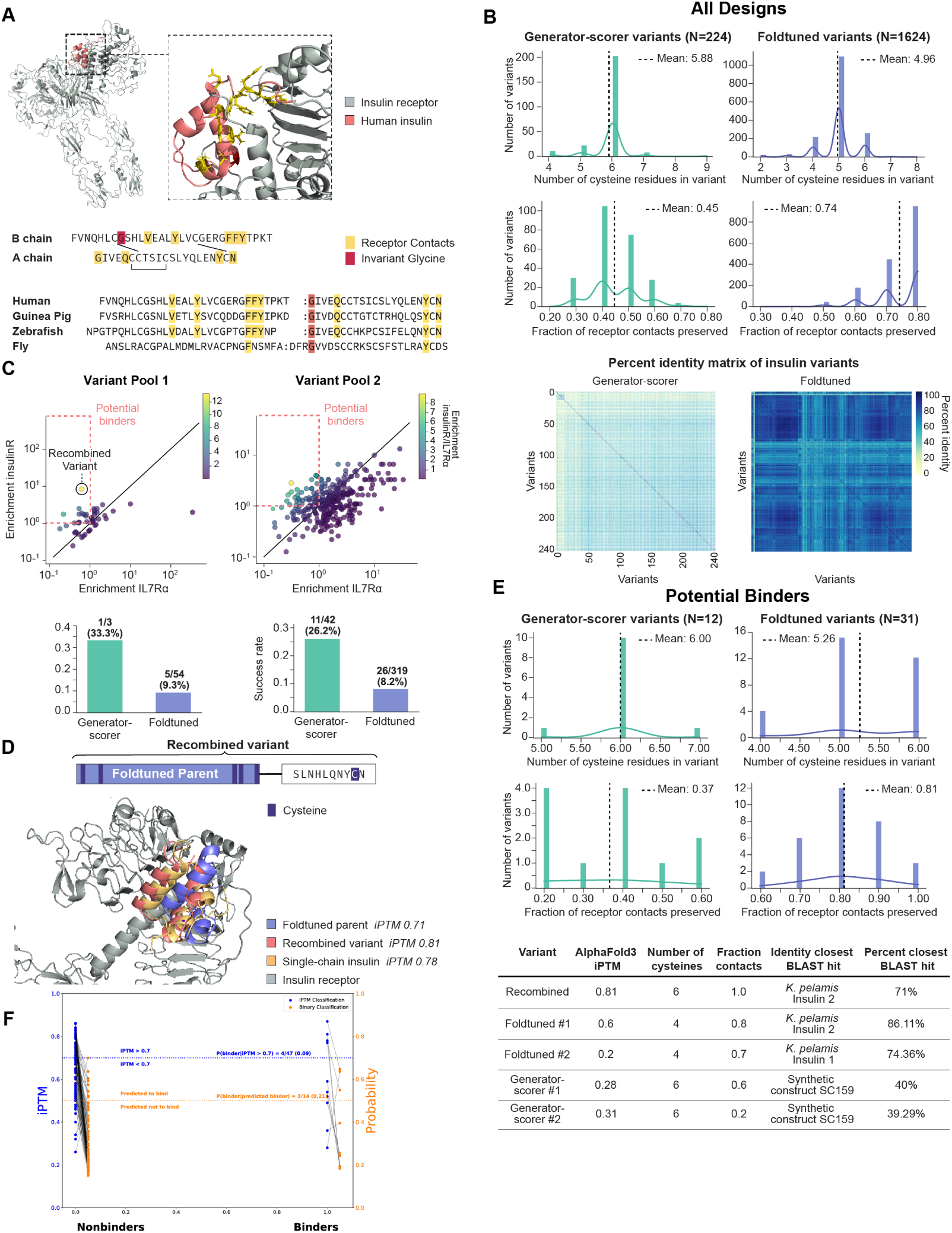
Protein CREATE illuminates properties of successful insulin receptor binder design. a) The human insulin sequences for the B-chain and A-chain are shown, linked via one intrachain and two interchain disulfide bridges. Residues implicated in binding to the human receptor are highlighted, along with an invariant glycine essential for proper folding. Sequence alignments of insulin homologs from other species are shown with preserved receptor contacts highlighted. The structure of human insulin with highlighted receptor contacts is also shown (PDB: 6SOF). **b)** Two different design strategies were used to generate predicted insulin receptor binders, showing wide sequence and property diversity. Foldtuning preserves more receptor contacts but produces many variants with an unpaired cysteine relative to other design strategies. **c)** The designs were assayed for insulin receptor binding using Protein CREATE. Due to potential misfolded variants that could show nonspecific binding, binders were determined by taking variants enriched on insulin receptor binding but de-enriched on an off-target receptor (IL7R*α*). **d)** A recombination event between two foldtuned variants was shown to be the most enriched variant in our first screen. Further analysis indicated that the addition of the C-terminal end shown improves predicted structural characteristics while restoring a possible disulfide bond. **e)** Binders from two of the tested design strategies (inverse folded variants were not prevalent in the pool and thus not shown) were analyzed to determine the relative importance of key insulin properties. The prevalence of designs with an odd number of cysteines decreased for both designs; however, interestingly, generator-scorer designs showed a slight decrease in receptor contacts on average, while foldtuned variants showed a slight increase. The top five hits, based on having the highest on-target insulin receptor enrichment over off-target IL7R*α* ratio, were chosen for individual analysis. With the exception of the recombined variant, all variants show relatively low iPTM scores, suggesting that using iPTM as a prescreen for binders may lead to false negatives. **f)** A held-out test set of variants from both design strategies and variant pools are classified into binders or nonbinders based on either iPTM scoring (blue) or a model trained on experimental data (orange). Variants above the blue and orange lines are predicted as binders by each respective method and nonbinders otherwise. True labels for all variants are given on the x-axis.

Given the diversity in natural solutions to the insulin receptor binding problem, we hypothesized that individual design strategies would be biased toward exploring only subsets of viable protein binders. We utilized two design strategies to generate binding candidates to human insulin receptor with structures templated on single-chain insulins and insulin-like peptides: foldtuning [20] and a strategy we term “generator-scorer.” Foldtuning leverages iterative refinement of a protein large language model to generate sequences that have folds similar to those of natural insulins and insulin-like peptides with decreased sequence identity to natural sequences. Our generator-scorer approach uses a protein large language model to propose sequences that are then evaluated by a scoring algorithm trained to predict Alphafold2 interchain predicted template modeling (iPTM) score between designs and the insulin receptor. Sequences that pass the scoring algorithm are then experimentally tested and used to update the generator model to produce more candidates to test.

As hypothesized, each design strategy produced sequences with different biochemical characteristics.

Foldtuned variants preserved most receptor contacts, but, interestingly, on average contain 5 cysteine residues as opposed to the 6 needed to form 3 disulfide bonds. The generator-scorer variants, by contrast, contain 6 cysteines like wild-type human insulin, but preserve far fewer receptor contacts (Figure 4b). As the reducing environment of *E. coli* cytoplasm presents a barrier to proper disulfide bond formation, we expected many phage-displayed variants would misfold, leading to phage dropout or nonspecific binding variants.

Leveraging the power of Protein CREATE to screen against multiple targets in parallel, we compared variant enrichment against Insulin receptor to that against IL7R*α* (Figure 4c) in two separate screens. As expected, many variants were enriched on both IL7R*α* and Insulin receptor, indicating nonspecific binding. Potential binders were identified by selecting variants with enrichment *>* 1 for Insulin receptor, but *<* 1 for IL7R*α*. We identified 44 putative binders using this filter (12 generator scorer and 31 foldtuned variants along with 1 variant that was the product of a recombination event).

The most enriched variant in our first screen was not created by either design strategy, but rather was the product of a recombination event between two foldtuned variants (Figure 4d). This recombined variant is predicted to have superior structural and biochemical properties compared to either foldtuned parent, as it contains no unpaired cysteines, preserves all insulin receptor contacts, and has a structure similar to that of single-chain human insulin when co-folded with the insulin receptor.

Of the other 43 variants selected from our filter, we see a strong selection in favor of variants without unpaired cysteines for both design strategies, while each design strategy converges on different ways to make contact with the insulin receptor. Foldtuned variants that preserve receptor contacts are enriched, while generator-scorer variants undergo no such selective pressure (Figure 4e). These trends are also observed when picking out individual variants most enriched in the assay. While all variants contain no unpaired cysteines, the variants differ markedly in the fraction of receptor contacts preserved. Interestingly, a majority of the enriched variants have an iPTM score less than 0.6, indicating that a subset of viable designs may be excluded from design campaigns that use Alphafold-based filtering as a pre-screen. Additionally, as hypothesized, key sequence features are conserved within design strategies but differ between them as evidenced by the identity of the closest BLAST result for each of the top enriched variants.

To demonstrate how Protein CREATE data can improve future rounds of design, we trained a binary classifier to predict insulin receptor binders from protein sequence using the data collected from our assay. We compared model performance on a held-out test set to predictions made using iPTM alone (Figure 4f). The model is able to reduce the high false positive rate of 91% (43/47) from iPTM scoring to 79% (11/14). This performance should be able to be further improved by collecting more data on binders due to the relative class imbalance currently seen in the data.

## Discussion

Experimental techniques for screening binder libraries have existed for almost 40 years [21]. Though this means ample time has passed for the development and optimization of screening technologies, it also means that these screening technologies were developed without regard for the promise - and limitations - of recent developments in generative AI-enabled protein design. For instance, traditional phage display relies on multiple rounds of selection and amplification in order to screen libraries against a desired target, which results in identification of only a handful of binders rather than quantification of each library member’s binding strength [22]. Phage display was also used to screen antibody libraries, identifying the first human antibody approved for therapy, Adalimumab (Humira®), and cDNA libraries rather than designed proteins generated by artificial intelligence algorithms [23]. Here, we create a novel experimental platform specialized for the mass collection of data to facilitate computational design of proteins.

Our results demonstrate the applicability of the platform on multiple targets. These targets can be multiplexed within a single assay, providing the dual advantage of cost and time savings while also quantifying off-target binding behavior. Furthermore, in contrast with traditional display technologies, Protein CREATE enables testing of multiple design strategies in addition to purely randomized libraries. Importantly, this control over library composition means that Protein CREATE can be used to test human or AI-generated hypotheses about design features at scale.

Limits on oligo pool lengths and variant dropout currently inhibit the applicability and throughput of Protein CREATE, respectively, but neither impose theoretical bounds to what should be possible with further refinement of the method. DNA assembly techniques such as Gibson and Golden Gate Assembly can be leveraged create larger protein constructs, while the T7 phage backbone used in Protein CREATE can accept exogenous DNA *>*3000 bp [22]. T7 phage titers can reach 10^10^ pfu/mL, meaning that 100 mL stocks can yield ∼ 1 trillion individual phage. Even assuming 1000 copies of each species, ∼ 1 billion variants could be screened in a single library. Recent progress in bacteriophage genome assembly and editing within an *E. coli* cell free lysate system may aid in surpassing current limits of the platform as well as reduce design-build-test cycle times [24].

As Protein CREATE is applied to more targets and collects more data, a generalized framework for incorporating the data received into a generative AI model will become more critical. We are exploring reinforcement learning paradigms, such as soft actor-critic [25], to encourage generative models to explore binders that are diverse in both sequence and structure space. Through continual collection and integration of data, we aim to create a “world model” of protein binding interactions, enabling fine-grained precision in designing novel protein binders for the bioengineering community.

## Acknowledgments

We thank Dr. Jost Vielmetter at the Caltech Protein Expression Center in the Beckman Institute for helping to perform and analyze SPR measurements. The authors also thank Inna Strazhnik for figure design and layout. This work was supported by Eli Lilly and Company through the Lilly Research Award Program (LRAP), the Gordon and Betty Moore Foundation, the Chan-Zuckerberg Initiative, the Beckman Foundation, and NIH R01-GM150125.

## Materials and Methods

### Phage Library Preparation

DNA Oligonucleotide Pools are ordered from Twist Biosciences and amplified for 12 cycles using Phusion High Fidelity PCR Master Mix (NEB, M0531S) in accordance with manufacturer’s instructions. This fragment is purified using CleanNGS beads (CleanNA, CNGS-0001) at a 0.8x sample ratio and washed twice with 80% ethanol before resuspension in ultra-pure water. The product is then assembled with T7Select 10-3b EcoRI/HindIII Vector Arms using NEbuilder HiFi Assembly Master Mix (NEB, E2621S) in accordance with manufacturer’s instructions. 5 µL of the reaction is added to 25 µL of T7Select packing kit (Millipore-Sigma, 70014-M) and incubated at room temperature for 2 hours. The entire reaction is directly added to top agar containing 10% (v/v) BLT5043 *E. coli* grown to mid-log phase and poured onto LB-agar containing plates. After incubation at 29ºC for 12 hours, plaques are lifted into the media using *Tris* − HCl, pH 8.0, 100 m**M** *NaCl*, 6m**M** *MgSO*_4_ and placed at 4ºC for at least 2 hours. The resulting supernatant is passed through a 0.2 µm filter and concentrated using 100 kDa MWCO filters (ThermoFisher Scientific, 88503). A buffer exchange with PBS is performed, followed by addition of 5x phage precipitation buffer (20% PEG 8000, 2.5 **M** NaCl). After incubation at 4ºC for 12 hours, the tube is centrifuged at 12,000 x g for at least 10 minutes and the precipitate resupended in 100 µL. The resultant phage solution is stored at 4ºC until further use.

### Binding Assay

For each binding assay, 0.3 mg of magnetic streptavidin beads (Acrobiosystems, SMB-B01-5mg) are washed 3 times with PBS before 1-2 µg of biotinylated target receptor is coupled to the bead by incubation for 1 hour at room temperature with agitation to keep the beads from settling. After 3 further washes with PBS, 30 µL of phage library is added to the beads and incubated at room temperature for 1-2 hours with agitation to keep the beads from settling. Beads are then washed ten times with PBST with 5.0% BSA.

A sample of the phage library and all of the washed beads are added to separate tubes and treated with DNAse I to digest unpacked phage genome followed by Proteinase K digestion of the phage capsid and the previously added DNAse. A single-cycle PCR is performed with a single UMI-containing primer using KAPA HiFi HotStart ReadyMix (Roche, KK2601) with the following thermocycling conditions: 98^º^C for 10 seconds, 50^º^C for 15 seconds, and 68ºC for 3 minutes. The PCR product is purified using CleanNGS beads (CleanNA, CNGS-0001) at a 0.8x sample ratio and washed twice with 80% ethanol before resuspension in ultra-pure water. Three dilutions (10x, 100x, 1000x) of each reaction product are made to determine a linear range in which UMI-based quantification is accurate. Each dilution and original sample is amplified using KAPA HiFi HotStart ReadyMix and primers to append Illumina P5 and P7 regions to the end of fragments. The PCR product is purified using CleanNGS beads (CleanNA, CNGS-0001) at a 0.8x sample ratio and washed twice with 80% ethanol before resuspension in ultra-pure water and then sequenced on an Illumina MiSeq NGS sequencer.

### Sequencing Data Analysis

We used a custom pipeline to analyze fastq output from the MiSeq sequencer. Briefly, we process sequences by removing sequences without the shared priming region, collapsing sequences containing UMI regions with Levenstein distance less than distance 3 away, deemed to be identical UMI regions, translating unique sequences into amino acid strings, and then matching these strings against a database of designed variants in the library. Each unique sequence, whether in the database or not, is then counted. We calculate enrichment using dilutions that have on average 10x coverage, as this is where UMI-based quantification is most accurate. Reads within the library before and after binding are normalized to have the same number of reads in each before enrichment is calculated as normalized unique reads after binding divided by normalized unique reads before binding for each species in the population.

### Molecular Dynamics Simulations

Molecular dynamics (MD) simulations were performed using GROMACS version 24.4 [26] with the AMBER99SB force field [27] and the TIP3P water model [28]. The initial structures of the receptorligand complexes were obtained from AlphaFold 3. The system was solvated in a dodecahedron box with water, and counterion Na+ was added to neutralize the system. Energy minimization was performed using the steepest descent algorithm until the maximum force was below 1000 kJ/mol/nm. The system was first equilibrated at 300 K for 100 ps using the modified Berendsen thermostat [29] in the NVT ensemble. The system was then equilibrated at 1 bar pressure for 100 ps using the Parrinello-Rahman barostat [30] in the NPT ensemble. The production MD was performed in the NPT ensemble at 300 K and 1 bar for 10 ns. All covalent bonds involving hydrogen atoms were constrained using the LINCS algorithm [31]. A cutoff of 1.0 nm was used for short-range electrostatic and van der Waals interactions. A timestep of 2 fs was used, with structures recorded every interval of 10 ps. Key properties including interaction energies, root-mean-square deviation (RMSD), and distance between receptor and ligand were analyzed using built-in GROMACS tools.

### Protein Expression and Purification

Linear fragments encoding each variant sequence, including a N-terminal leader sequence consisting of MSHHHHHHHHSENLYFQSGGG, along with a strong promoter, ribosome binding site, and terminator were ordered from Twist Biosciences as double-stranded linear DNA fragments. This DNA was amplified using PrimeStar GxL DNA Polymerase (Takara, R051A) in accordance with manufacturers instructions and purified using CleanNGS beads (CleanNA, CNGS-0001) at a 0.8x sample ratio and washed twice with 80% ethanol before resuspension in ultra-pure water. 15 µL of DNA was added to 45 µL of myTXTL Pro Cell-Free Expression Master Mix (Arbor Biosciences, 540300) and incubated at 29ºC for 12 hours. The resultant protein product was purified using Ni-NTA affinity chromatography (NEB, S1427) in accordance with manufacturer’s instructions. Following purification, eluted protein is buffer exchanged into PBS and concentrated using a 3k kDa MWCO filter (Millipore Sigma, UFC5003) and stored at 4^º^C until further use.

### Surface Plasmon Resonance (SPR) Measurements

SPR was performed using the Bruker SPR-32 system using HBS-EP+ buffer (10 m**M** HEPES pH 7.4, 150 m**M** NaCl, 3 m**M** EDTA, 0.005% v/v) Surfactant P20, GE Healthcare). Biotinlyated IL7R*α* (Acrobiosystems, IL7-H82F9-25ug) was immobilized on Biotin Tag Capture Sensor chips (Bruker, 1862620) until 600 relative units were immobilized. Increasing concentrations of IL7R*α* binders were injected at a flow rate of 30 µL per minute. Concentrations tested for each binder were 781 p**M**, 1.56 n**M**, 3.125 n**M**, 6.25 n**M**, 12.50 n**M**, 25.0 n**M**, and 50.0 n**M**. Curves were fit and a *K*_*D*_ derived using the titration cycle kinetic model within Bruker R4 analysis software.

### Luminex Immunodetection Assay

Multiple sets of Bio-Plex magnetic COOH beads (Bio Rad, 171406001) were coupled with streptavidin or avidin in accordance with manufacturer’s instructions. Post-coupling, beads were incubated with either biotinlyated IL7R*α* (Acrobiosystems, IL7-H82F9) or IL2R*βγ* (Acrobiosystems, ILG-H82F3) in blocking buffer (PBST + 0.01% thiomersal and 2.5% BSA) to immobilize biotinylated receptor on beads for 1 hour. Beads were then blocked with 1 mM biotin, washed, and incubated with 1 µ**M** of His-tagged binder as laid out in Figure 2d for 30 minutes. The beads were stained for 30 minutes with 6x His tag monoclonal antibody from mouse (Genscript, A00186), followed by a wash and another 30 minute incubation with secondary stain with phycoerythrin-labeled anti-mouse IgG antibody from goat (Biolegend, 405307). Following a wash with PBS, the beads were analyzed using the Bio-Plex 200 system (Biorad).

### IL7R*α* Inhibitor Assay

HEK-Blue™ IL-7 Cells were purchased from Invivogen and passaged 7 times in DMEM containing 4.5 g/l glucose and 2 m**M** L-glutamine, 10% (v/v) heat-inactivated fetal bovine serum, 100 U/ml penicillin, and 100 *µ*g/ml streptomycin. At the time of the assay, after a fresh change of media, approximately 50,000 cells in 180 *µ*L of media were added to a 96-well plate. Varying concentrations of each inhibitor as described in the text were added and the cells incubated for 2 hours at 37^º^C 5% *CO*_2_. After incubation, human IL-7 was added to each well to a final concentration of 17 p**M** and the cells incubated for another 12 hours at 37^º^C 5% *CO*_2_. Secreted embryonic alkaline phosphatase (SEAP) was detected by measuring absorbance at 620 nm using QUANTI-Blue^™^ Solution (Invivogen) in accordance with manufacturer’s instructions after 45 minutes of incubation at 37ºC. Fractional response was calculated by bounding the average response of the cells to IL7 without either inhibitor at 1 and the average response of the cells without IL7 added at 0.

